# Detecting overlapping spatial clusters of high sugar-sweetened beverage intake and high body mass index in a general population: a cross-sectional study

**DOI:** 10.1101/399584

**Authors:** Stéphane Joost, David De Ridder, Pedro Marques-Vidal, Beatrice Bacchilega, Jean-Marc Theler, Jean-Michel Gaspoz, Idris Guessous

**Affiliations:** Laboratory of Geographic Information Systems (LASIG), School of Architecture, Civil and Environnemental Engineering (ENAC), Ecole Polytechnique Fédérale de Lausanne (EPFL), Switzerland; Unit of Population Epidemiology, Division of Primary Care Medicine, Department of Community Medicine, Primary Care, and Emergency Medicine, Geneva University Hospitals; Faculty of Medicine, University of Geneva, Switzerland; Group of Geographic Information Research and Analysis in Population Health (GIRAPH); Department of Medicine, Internal Medicine, Lausanne University Hospital, Switzerland; Institute of Social and Preventive Medicine (IUMSP), Division of chronic diseases (dMC), Lausanne University Hospital (CHUV), Lausanne, Switzerland

**Keywords:** sugar-sweetened beverage, body mass index, spatial dependence, clusters

## Abstract

**Objective:** To identify populations and areas presenting higher consumption of sugar-sweetened beverages (SSB) and their overlap with populations and areas presenting higher body mass index (BMI).

**Design:** Cross-sectional population-based study.

**Setting:** State of Geneva, Switzerland.

**Participants:** 15,767 non-institutionalized residents aged between 35 and 74 years (20 and 74 since 2011) of the state of Geneva, Switzerland.

**Main outcome measures:** Spatial indices of sugar-sweetened beverage intake frequency and body mass index. Median regression analysis was used to control for characteristics of patients.

**Results:** The SSB intake frequency and the BMI were not randomly distributed across the state. Among the 15,423 participants retained for the analyses, 2,034 (13.2%) were within clusters of high SSB intake frequency and 1,651 (10.7%) was within clusters of low SSB intake frequency, 11,738 (76.1%) showed no spatial dependence. We also identified clusters of BMI, 4,014 (26.0%) participants were within clusters of high BMI and 3,591 (23.3%) were within clusters of low BMI, 7,818 (50.7%) showed no spatial dependence. We found that clusters of SSB intake frequency and BMI overlap in specific areas. 1,719 (11.1%) participants were within high SSB intake frequency and high BMI clusters. After adjustment for covariates (education level, gender, age, nationality, and the median income of the area), the identified clusters persisted and were only slightly attenuated.

**Conclusion:** A fine-scale spatial approach allows identifying specific populations and areas presenting higher SSB consumption and, for some areas, higher SSB consumption associated with higher BMI. These findings could guide legislators to develop targeted interventions such as prevention campaigns and pave the way for precision public health.

**What is already known on this topic:**

- The consumption of sugar-sweetened beverages (SSBs) is an important contributory factor of obesity and obesity-related diseases.
- SSB consumption varies according to socioeconomic status, which could explain the higher prevalence of obesity in specific areas.
- SSB taxation faces resistance in many countries due to its potential regressive nature.

**What this study adds:**

- The spatial analysis of individual-level SSB consumption in the state of Geneva provides a clear identification of populations and areas presenting higher SSB consumption and, for some areas, higher SSB consumption along with higher body mass index (BMI).
- The results demonstrate the persistence of SSB clustering in the geographic space after adjusting for education level, gender, nationality, age, and neighborhood-level median income.
- The findings provide guidance for future public health interventions to reduce SSB consumption by better targeting vulnerable populations.

## Introduction

The prevalence of obesity and obesity-related diseases has been increasing steadily in most countries over the past decades (1). Although pathways leading to obesity are multiple and complex, notably because of the interplay of genetic, environmental, and social factors, it has been suggested that one important contributory factor is the consumption of sugar-sweetened beverages (SSB) (2,3). Sugar-sweetened beverages are drinks with added sugar and include a wide range of products such as soft drinks, flavoured juice drinks, sports drinks, sweetened tea, coffee drinks, energy drinks, and electrolyte replacement drinks (4). While these beverages present significant differences in sugar content, drinks such as sodas can contain up to 39 g of sugar per 330 ml can (5). Worldwide, SSB sales and consumption have increased these last decades with the greatest consumptions reported in Argentina and in the United States (6). In Europe, the annual *per capita* consumption of SSB was about 95 litres in 2015 (7). In Switzerland, SSB consumption increased during the last 20 years among children, teenagers, and adults to reach an annual consumption of about 80 litres (8).

Hence, SSB have a growing contribution to total energy intake and may lead to obesity and obesity-related diseases including diabetes, hypertension, and cardiovascular diseases (9). In addition to the increase in energy intake, SSB may also impact health via the metabolic response to fructose, a major component of SSB. High intake of fructose can lead to increased visceral adiposity, lipid dysregulation, and decreased insulin sensitivity (10). Several studies have suggested a relationship between excessive consumption of SSB and obesity; although some controversy remains on whether the association is causal, a recent systematic review and meta-analysis of large prospective cohort studies and randomized controlled trials concluded that SSB consumption promotes weight gain in children and adults (11). Recent meta-analyses also concluded that usual consumption of SSB was associated with a greater risk of hypertension (12,13), stroke (14,15), and type 2 diabetes, independently of adiposity (16). Accordingly, several governmental and public health interventions have been implemented to reduce consumption of SSB (17).

SSB taxation, already introduced in some states in the USA and in several other countries, has a great potential to reduce SSB consumption. However, objections against such tax have raised in many countries, notably due to its regressive nature and supposed lack of efficacity to lower obesity prevalence (18). Nevertheless, it has been suggested that these arguments could be addressed by ensuring that the revenues generated are allocated preferentially to programs promoting nutrition and obesity-prevention for the most in need (18). Therefore, identifying specific populations at risk of SSB overconsumption is of utmost importance for health policymakers. Still, the identification of such populations or areas in need of intervention is far from optimal.

Spatial analysis methods have been developed and introduced in epidemiological research to explore the link between place of residence and health (19). Spatial clusters of a specific trait can be detected by its spatial dependence (spatial autocorrelation), defined as the covariation of properties in the geographic space (20,21).

Here, we propose a fine-scale geographical approach to identify populations and areas in high need for interventions to reduce SSB consumption. Several studies explored the spatial distribution of SSB at the county or state-level (22–24), what results in a smoothing and altering the original signal by aggregating individual-level data. The second aim was to identify areas where clusters of high SSB intake frequency overlapped with clusters of high BMI. Such areas would be ideal targets for intervention, eventually supported by revenue generated from SSB taxation.

## Methods

### Data source and study population

Data on adults were collected using the Bus Santé study (25), a cross-sectional population-based study that collects information on cardiovascular risk factors. Every year, a stratified sample of 500 men and 500 women representative of the State of Geneva’s 100’000 males and 100’000 females non-institutionalized residents aged 35–74 (20-74 since 2011) is recruited and studied. Four trained collaborators interview and examine the participants. All procedures are reviewed and standardized across technicians on a regular basis.

Eligible subjects are identified via a standardized procedure using an annual residential list established by the local government. This list includes all individuals living in the State of Geneva. An invitation to participate is mailed to the sampled subjects and, if they do not respond, up to seven telephone calls are made at different times on various days of the week. If telephone contact is unsuccessful, two more invitation letters are sent. Subjects that are not reached are replaced using the same selection protocol. Subjects who refuse to participate are not replaced. Subjects who accept to participate receive a self-administered, standardized questionnaire including a semi-quantitative food frequency section (FFQ). Geographic coordinates of the postal address are used for individual geographic information. For this analysis, data from surveys 1995 to 2014 were used, corresponding to 15,767 participants. The average participation rate for 1995-2014 was 61% (range: 53–69%).

### Body Mass Index and Sugar-Sweetened Beverages Intake Frequency

Participants brought filled-in questionnaires, which were checked for correct completion by trained interviewers (26). Body weight was measured with the subject lightly dressed, without shoes and using a medical scale (precision 0.5□ kg); standing height was measured using a medical gauge (precision 1□cm). Body mass index (BMI) was calculated as weight (kg)/height (m^2^).

SSB intake frequency was assessed for every participant using a self-administered, semiquantitative food frequency questionnaire (FFQ), which also included portion sizes (27,28). This FFQ has been validated against 24-h recalls among 626 volunteers from the Geneva population, and data derived from this FFQ have recently contributed to worldwide analyses (29). Briefly, this FFQ assesses the dietary intake of the previous 4 weeks and consists of 97 different food and beverage items, including SSB (colas, sodas, lemonades, syrups). For each item, consumption frequencies ranging from “less than once during the last 4 weeks” to “2 or more times per day” were provided; daily SSB intake frequency was computed from 0 for “less than once during the last 4 weeks” to 2.5 for “2 or more times per day”. Information derived from this FFQ has contributed to several reports from large consortium such as the Global Burden of Disease (30).

The local Institutional Ethics Committee approved the study. All participants gave a written informed consent prior to entering the study.

### Covariates

SSB consumption and BMI covariates included education level, gender, age, nationality, and the neighborhood-level median income of the area. We used income data characterizing the 475 Geneva statistical sectors in 2009 for adjustment. These data were produced by *Statistique Genève* (31). The yearly income value (1 CHF = 1.007 USD, June 2018) was attributed to Bus santé participants based on their postal address within the corresponding statistical sectors.

### Statistical analyses

Using the geographical coordinates of the place of residence, we used the Getis-Ord Gi statistic (32,33) implemented in the Geoda software (34) to investigate whether SSB intake frequency and BMI were spatially dependent. Getis-Ord Gi indicators are statistics that measure spatial dependence and evaluate the existence of local clusters - hot or cold spots - in the spatial arrangement of a given variable, here SSB intake frequency and BMI. They compare the sum of individual’s SSB intake frequency values included within a given spatial lag proportionally to the sum of individual’s SSB intake frequency values within the whole study area, and similarly for BMI (32). The Gi statistic returned for each value is a Z-score (standardized value) to which a p-value is associated. The null hypothesis for this statistic is that the values being analyzed exhibit a random spatial pattern. When the p-values are statistically significant, it can be assumed that the spatial distribution is not random. Statistical significance testing was based on a conditional randomization procedure (35) using a sample of 999 permutations. Large statistically significant positive Z-scores reveal clustering of high values, while large significant negative Z-scores reveal clustering of low values.

We decided to show the results of the analysis of SSB intake frequency and BMI variables using a spatial lag of 1,200m around the place of residence of each individual as this distance approximates the size of a typical neighborhood in the urban areas of the studied territory. No threshold could be determined on the basis of a quantitative criterion: a correlogram calculated with a maximum spatial lag of 4km produced global Moran’s I ranging between 0 and 0.011 for BMI in the ten 400m-bins, and between 0 and 0.001 for SSB intake frequency. Considering a correlogram calculated with a maximum distance of 2km, Moran’s I for BMI ranged between 0.002 and 0.016 in the ten 200m-bins, and between 0 and 0.003 for SSB intake frequency, translating no global spatial autocorrelation in both variables.

We used a standardized approach where the sum of the weights (W) equals 1 and each individual weight is 1/W_i_. In this case, Gi and Gi* (including the value of the target individual) are homogeneous of order zero in W_ij_ and thus invariant (33).

We used a median regression analysis to obtain the SSB intake frequency and the BMI adjusted for education level, gender, age, nationality, and the median income of the area (36). Statistical significance was considered for a p-value < 0.05 for all spatial dependence measures.

Finally, to determine whether SSB intake frequency, BMI and their spatial dependence were stable during the 1995-2014 period, we divided the dataset into 3 subperiods with a high number of participants to favor the robustness of the evaluation – P1 = 1995-2001 (n = 5,511), P2 = 2002-2008 (n = 4,714), and P3 = 2009-2014 (n = 5,357) – and processed for each of them i) Tukey’s multiple comparison analysis method to ensure that the mean of the SSB-IF and of the BMI between the 3 periods had not increased or decreased sharply and ii) Moran’s I to verify that there was no difference in global spatial autocorrelation between the 3 periods (37).

On the maps produced, white dots show locations where there is no spatial dependence. Red dots show individuals with a statistically significant positive Z-score, meaning that high values cluster within a spatial lag of 1,200 m, and are found closer together than expected if the underlying spatial process was random. Blue dots show individuals with a statistically significant negative Z-score, meaning that low values cluster within a spatial lag of 1,200 m, and are found closer together than expected if the underlying spatial process was random. We presented maps derived from unadjusted and area’s income-adjusted SSB intake frequency or BMI, respectively.

In order to compare the possible overlap between SSB intake frequency and BMI spatial clusters, participants were categorized in 10 classes (**Table S1**) using the combination of the previously computed Getis-Ord Gi clusters SSB intake frequency and BMI. The same classification was performed before and after adjustment for covariates.

## Results

After excluding participants for missing data, 15,423 (97.8%) participants were retained. Genders were represented at the same rate (50.0%), the mean age of the participants was 51.3 years (SD ± 11.0 years), 37.7% of the participants had a university level degree, 70.6% were of Swiss nationality and 29.4% of other nationalities, the neighborhood-level median yearly income was 72,166 CHF. The mean SSB intake frequency was about 0.22 SSB/day (SD ± 0.5 SSB/day) and 0.18 SSB/day (SD ± 0.5 SSB/day) after adjustment for covariates. Only 3.5% of participants consumed SSB twice or more per day and 5.8% once a day. The mean BMI was 24.9 kg/m^2^ (SD ± 4.1 kg/m^2^) and 25.0 kg/m^2^ (SD ± 3.8 kg/m^2^) after adjustment for covariates. Around half of participants had a normal weight (49.5%, BMI between 18.5 and 25 kg/m^2^) and 10.6% of participants had a BMI ≥ 30 kg/m^2^.

Analyses of mean and trends of BMI and SSB intake frequency (**Table S2**) as well as Moran’s I (**Table S3**) across the 3 subperiods are presented in Supplementary material. Despite a slight increase of BMI and SSB intake frequency over time, the difference was only significant between P1 (1995-2001) and P3 (2009-2014) for both raw and adjusted variables (**Figure S1**). The absence of global spatial autocorrelation for both variables was stable during the three periods while the spatial distribution of local clusters of BMI and SSB intake frequency slightly vary (SSB intake frequency hotspot downtown during P1 only) (**Figure S2A-C**).

### Sugar-Sweetened Beverages Intake Frequency clusters

Before adjustment, 11,738 individuals (76.1%) showed no spatial dependence, whereas 2,034 (13.2%) had a high SSB intake frequency (constituting hot spots) and 1,651 (10.7%) a low SSB intake frequency (cold spots) (**Figure 1A**). After adjustment, 11,936 individuals (77.4%) showed no spatial dependence, 2,011 (13.0%) were included within SSB intake frequency hot spots and 1,476 (9.6%) in SSB intake frequency cold spots (**Figure 1B**). Both maps show a clear global structure with SSB intake frequency cold spots mainly located to the east of the lake (see landmark #6 on maps) and SSB intake frequency hot spots to the west (#1, 3, 4, 9). The main effect of adjustment is to attenuate the geographic footprint of hot and cold spots.

**Figure 1:**
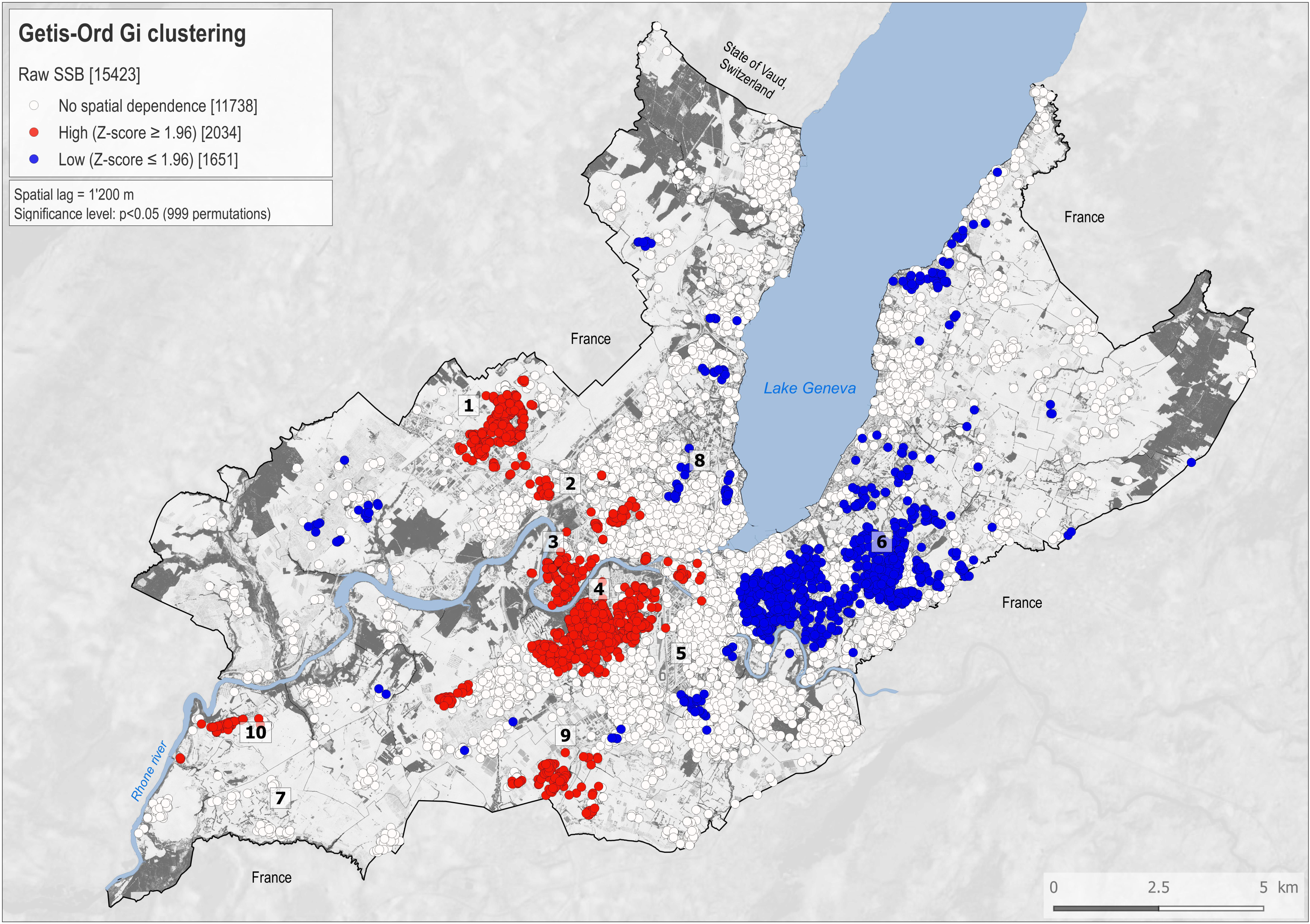
Getis-Ord Gi clusters calculated for 15,423 Bus santé participants (1995-2014) for the raw sugar-sweetened beverage (SSB) intake frequency variable (A) and adjusted for covariates (B). White dots show places where the space is neutral (no spatial dependence). Red dots show individuals with a statistically significant positive Z-score (α=0.05), meaning that high values cluster within a spatial lag of 1,200m and are found closer together than expected if the underlying spatial process was random. Blue dots show individuals with a statistically significant negative Z-score (α=-0.05), meaning that low values cluster within a spatial lag of 1,200m and are found closer together than expected if the underlying spatial process was random. Indicative landmarks numbered 1-10 are displayed on the maps and used to support the description of the results.

### BMI clusters

Before adjustment, 7818 participants (50.7%) showed no spatial dependence, 4,014 (26%) were located within BMI hot spots and 3,591 (23.3%) within BMI cold spots (**Figure 2A**). After adjustment, 8,253 participants (53,5%) showed no spatial dependence, 3,409 (22.1%) were located within BMI hot spots and 3,761 (24.4%) within BMI cold spots (**Figure 2B**). The effect of adjustment thins down the large hot spot located between #1 and #5 and reduces and shifts the large cold spot located between #5 and #6 towards the west.

**Figure 2:**
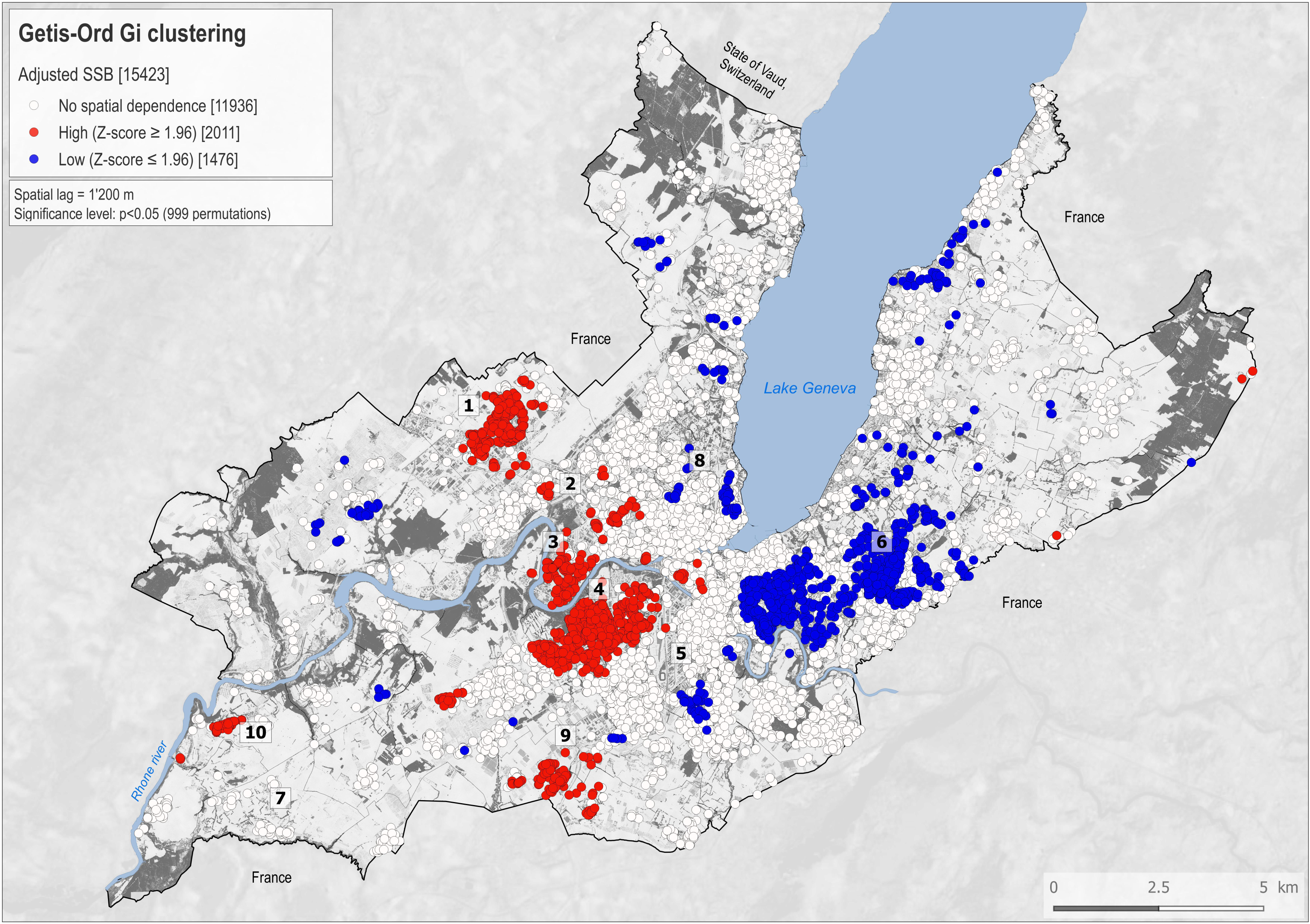
Map showing Getis-Ord Gi clusters calculated for 15,423 Bus santé participants (1995-2014) for the raw body mass index (BMI) variable (A) and adjusted for covariates (B). White dots show places where the space is neutral (no spatial dependence). Red dots show individuals with a statistically significant positive Z-score (α=0.05), meaning that high values cluster within a spatial lag of 1,200m and are found closer together than expected if the underlying spatial process was random. Blue dots show individuals with a statistically significant negative Z-score (α=-0.05), meaning that low values cluster within a spatial lag of 1,200m and are found closer together than expected if the underlying spatial process was random. Indicative landmarks numbered 1-10 are displayed on the maps and used to support the description of the results.

### Spatial overlap between SSB intake frequency and BMI clusters

A spatial overlap between SSB intake frequency and BMI clusters (i.e., co-location of hot spot SSB intake frequency and hot spot BMI - class 1) was clearly identified and included 1,719 (11.1%) participants (**Figure 3A**). After adjustment for education level, gender, age,nationality and the median income of the area, overlapping clusters included 1,595 (10.3%) participants (**Figure 3B**).

**Figure 3:**
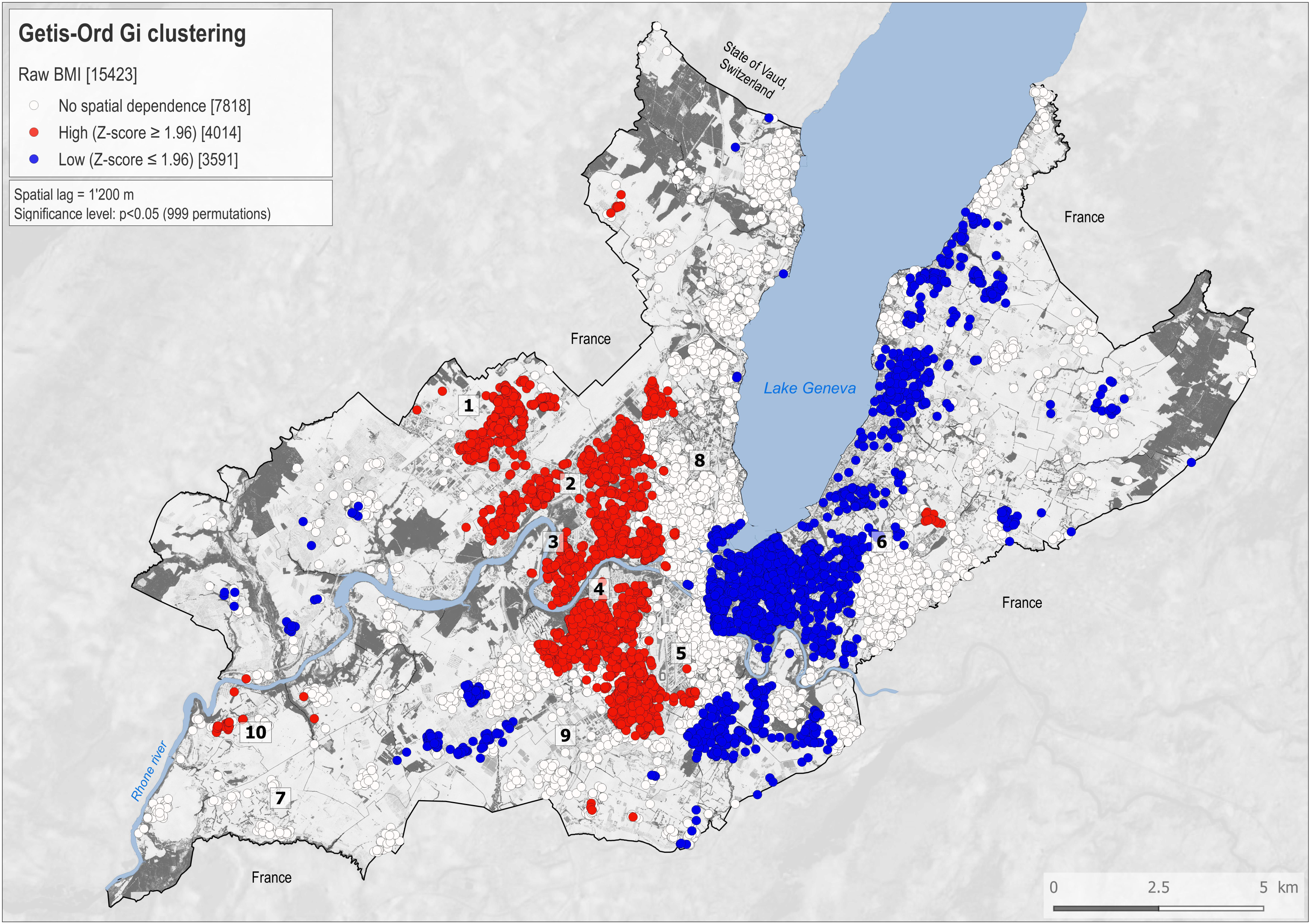
The main delimited clusters with individuals belonging to both raw SSB intake and raw BMI hotspots contain 1,719 individuals. (A). The main delimited clusters with individuals belonging to the adjusted SSB intake frequency and BMI hotspots contain 1,595 individuals. (B). Indicative landmarks numbered 1-10 are displayed on the maps and used to support the description of the results.

## Discussion

To the best of our knowledge, this is the first study to report simultaneously SSB intake frequency and BMI spatial clustering in a general adult population. Using geo-referenced measurements of SSB intake frequency at the individual level, we identified significant clusters of high and low SSB intake frequency values, together with their spatial distribution in the State of Geneva. One eighth of the population was within clusters of high SSB intake frequency. We then identified clusters of BMI and found that clusters of SSB intake frequency and BMI overlapped in specific areas. One tenth of the population was simultaneously within high SSB intake frequency and high BMI clusters.

### Sugar-Sweetened Beverages Intake Frequency clusters

Compelling evidences suggest that SSB consumption increases the risk of obesity and obesity-related diseases such as diabetes and hypertension (11–13,16). The negative impact of SSB consumption on population health is expected to increase given the worldwide increasing trends in SSB consumption (38). Experts and organizations have advocated and conducted numerous interventions (17) aiming to prevent SSB overconsumption and its related negative health impact by increasing public awareness and change nutritional behavior (10). However, the identification of the most vulnerable populations in high need of interventions, enabled by spatial analysis methods, could improve the efficacy and efficiency of local level programs. These methods could be of even greater interest when considering SSB taxation as the effect of such dissuasion measures could be monitored in time. This approach has already been implemented in some regions, including the USA (18,39). Taxing SSB is an effective global public health approach to reduce SSB consumption. Still, many have expressed concerns against the regressive nature of the tax (18) contributing to the resistance of legislators and other stakeholders in implementing such largescale public health policy in other regions. To overcome this barrier, it has been proposed to allocate the revenue generated from SSB taxation to implement nutrition programs or obesity-prevention programs. Ideally, these SSB tax-funded programs should preferentially target populations most vulnerable to SSB consumption. The identification of such populations or areas in need of intervention is currently challenging and would highly benefit from the contribution of local scale spatial approaches. Clusters of higher SSB intake frequency (and respectively lower SSB intake frequency) revealed by our analysis in specific areas of Geneva could constitute initial targets in such a precision public health context. Some of these areas have a lower socio-economic status than in other districts of Geneva, an urban state where recent evidence highlighted the existence of social inequalities in dietary intake (40). Neighborhood socio-economic status (e.g., neighborhood deprivation, neighborhood segregation, population density) is known to be a determinant of dietary habits, obesity, obesity-related diseases and even mortality (41). However, we also identified high SSB consumption clusters that were independent of education level, gender, age, nationality, and neighborhood-level median income suggesting that other factors such as network phenomena (e.g., social networks) (42) and environmental factors (e.g., types of food stores, food access) (43) influenced SSB intake frequency.

Other studies explored the spatial distributions of SSB at broader geographic scales, usually county or state-level (22–24). Indeed, the distribution of local-scale phenomena, such as SSB consumption, can be smoothed out by aggregating information to large administrative spatial units (e.g. county) and altering the original signal. Here, we did not used predefined unit, but instead considered space as a continuum. In 2017, Tamura et al., reported the spatial clustering of SSB consumption in adolescents using a sample of 1,292 precisely georeferenced residential address from the Boston youth study. They identified clusters with high rates of non-soda SSB (44). However, in their study, spatial clusters of SSB consumption did not resist adjustment for gender, education, age and ethnicity, suggesting that in Boston these factors might play a larger role in the determination of the spatial distribution of SSB consumption.

To provide even more specific information on where to implement programs and interventions on SSB and SSB-related health risk, we provided information on areas corresponding to overlapping clusters of higher SSB intake frequency and higher BMI. In line with our previous work (20), we found spatial clustering of BMI in adults from the general population and compared them to SSB intake frequency clusters. These overlapping areas could be interpreted as areas where individuals are already suffering the negative impact of SSB intake frequency on weight and potentially other related diseases. This information could further guide on the priority (high needed areas) and nature (programs focusing on the negative impact of SSB consumption on obesity) of the public health interventions.

In addition to guiding interventions, characterizing SSB intake frequency clusters and overlapping clusters at both the individual and environmental levels could be used for research purposes to further our understanding on the social and environmental determinants and consequences of SSB consumption. For example, the density of advertising (SSB or fast food advertisements) and SSB vending machine in the hot spot clusters could be determined as they have been associated with SSB consumption (43,45,46). This research effort could provide further evidence on the causality of sugar overconsumption on health consequences that remains controversial (47).

Our study is not without limitations. Firstly, regarding spatial statistic parameters, we chose to use a spatial lag of 1,200m, but other choices may produce slightly different results. Yet, we tested the robustness of our findings using different spatial lags and found no meaningful difference in the clusters obtained. Secondly, we favored the Getis-Ord Gi statistic instead of Local Indicators of Spatial Association (35) (LISA) as we focused primarily on the detection of local clusters of high values, keeping discordant behaviors for further future investigations. Thirdly, participants and non-participants in the Bus Santé study may differ regarding SSB consumption and participation bias cannot be excluded. Still, to reduce participation bias, the Bus Santé study has a mobile examination unit that covers three major areas of the State favoring the participation of people living in lower SES areas. Fourthly, SSB intake frequency was self-reported, and recall as well as social desirability biases cannot be excluded. Finally, preliminary analysis of 3 temporal groups (1995-2001; 2002-2008; 2009-2014) of SSB intake frequency and BMI showed a similar overall spatial structure as translated by a stable Moran’s I over time; local Getis Gi clusters are stable also, with the only exception of SSB intake frequency for P1 showing a slightly different pattern. The overall stability described above allowed us to perform an overall analysis over 20 years of population-based data. Finally, such an approach is easily applicable elsewhere as the variables used are frequently collected in medical cohorts. Their acquisition is also inexpensive. One difficulty, however, lies in being able to benefit from specific geographical data that precisely locate the building of residence of the participants.

## Conclusions and policy implications

Numerous programs and interventions have been conducted to mitigate obesity prevalence and SSB consumption. Yet, far more progress could be achieved by implementing interventions targeting the most vulnerable populations and creating a tax on SSB. Our investigation, founded on individual-level data, provides a fine scale assessment of the spatial clustering in SSB consumption, an important factor known to favor obesity. The identification of specific areas presenting higher SSB consumption and, for some specific areas associated with higher BMI values, enables local legislators and public health experts to develop targeted interventions and paves the way for precision public health delivery. Finally, our findings permit to allocate resources to the populations in high need of interventions, which could diminish resistance against SSB taxation.

### Detecting overlapping spatial clusters of high sugar-sweetened beverage intake and high body mass index in a general population: a cross-sectional study

Stéphane Joost^1,2,4,6*^, David De Ridder^1,2,3,4*^, Pedro Marques-Vidal^4,5^, Beatrice Bacchilega^1^, Jean-Marc Theler^2^, Jean-Michel Gaspoz^2,3^, Idris Guessous^2,3,4*^

## SUPPLEMENTARY MATERIAL

**Supplementary Table T1:** Classes resulting from the combination of Getis-Ord Gi clusters for SSB intake frequency and BMI.

| Classes | Description |
| --- | --- |
| 1 | SSB hot spot - BMI hot spot spot |
| 2 | SSB hot spot - BMI cold spot |
| 3 | SSB cold spot - BMI hot spot |
| 4 | SSB cold spot - BMI cold spot |
| 5 | SSB hot spot - BMI neutral |
| 6 | SSB cold spot - BMI neutral |
| 7 | SSB - BMI hot spot |
| 8 | SSB - BMI cold spot |
| 9 | Permanent neutral |

**Supplementary Table S2.** Trends of mean body mass index (BMI) and sugar-sweetened beverage (SSB) intake frequency across three subperiods P1 (1995-2001), P2 (2002-2008), and P3 (2009-2014).

|  | P1 | P2 | P3 |
| --- | --- | --- | --- |
| BMI (kg/m<sup>2</sup>) | 24.70 | 24.89 | 25.07 |
| BMI adjusted (kg/m<sup>2</sup>) | 24.69 | 24.90 | 25.07 |
| SSB (intake frequency per day) | 0.20 | 0.22 | 0.24 |
| SSB adjusted (intake frequency per day) | 0.20 | 0.22 | 0.24 |

The division of the global dataset into 3 subperiods (P1-P3) showed a slight increase of BMI and SSB intake frequency overtime, the difference is only significant between P1 and P3 for both the raw and adjusted variables (using the Tukey Multiple Comparison of Means, honest significance difference with a family-wise error rate equal to 0.05) (Figure S1).

**Supplementary Table S3.** Trends of spatial autocorrelation calculated with spatial statistic Moran’s I across three subperiods P1 (1995-2001), P2 (2002-2008), and P3 (2009-2014) for the variables BMI and SSB intake frequency. Statistical significance assessment using a conditional randomization procedure using a sample of 999 permutations with α= 0.05.

|  | P1 | P2 | P3 |
| --- | --- | --- | --- |
| BMI (kg/m<sup>2</sup>) | 0.0059** | 0.0114*** | 0.0100*** |
| BMI adjusted (kg/m<sup>2</sup>) | 0.0026 | 0.0103*** | 0.0071*** |
| SSB (intake frequency per day) | -0.0001 | 0.0002 | 0.0004 |
| SSB adjusted (intake frequency per day) | 0.0002 | 0.0004 | -0.0005 |

The absence of global spatial autocorrelation for both variables is stable during the three periods while the spatial distribution of local clusters of BMI and SSB intake frequency slightly vary (SSB intake frequency hotspot downtown during P1 only) (**Figure S2A-C**).

**Supplementary Figures**

**Figure S1:**
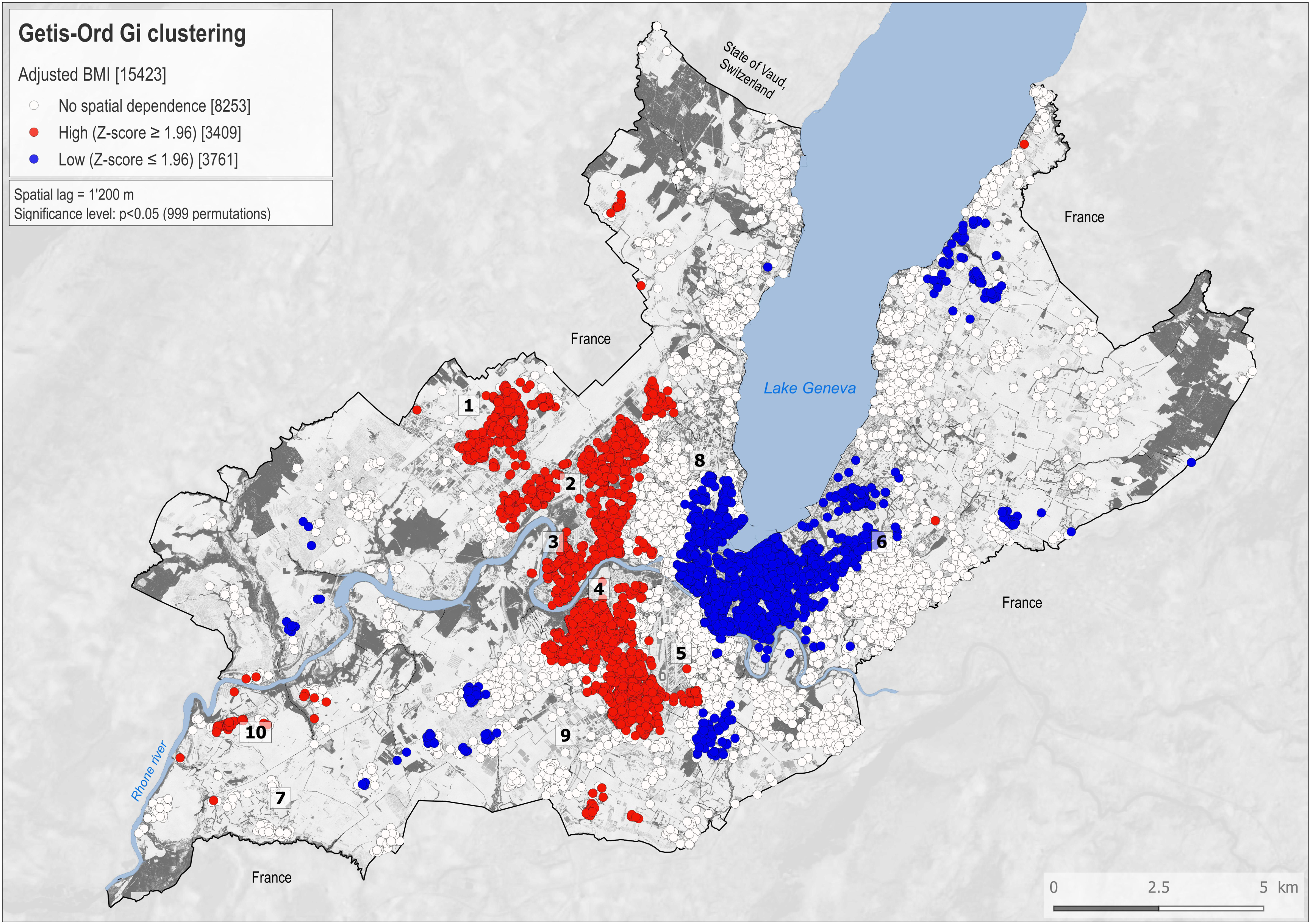
Mean BMI (A), adjusted BMI (B), SSB intake frequency (C), and adjusted SSB intake frequency (D) across subperiods P1 (1995-2001), P2 (2002-2008) and P3 (2009-2014)).

**Figure S2:**
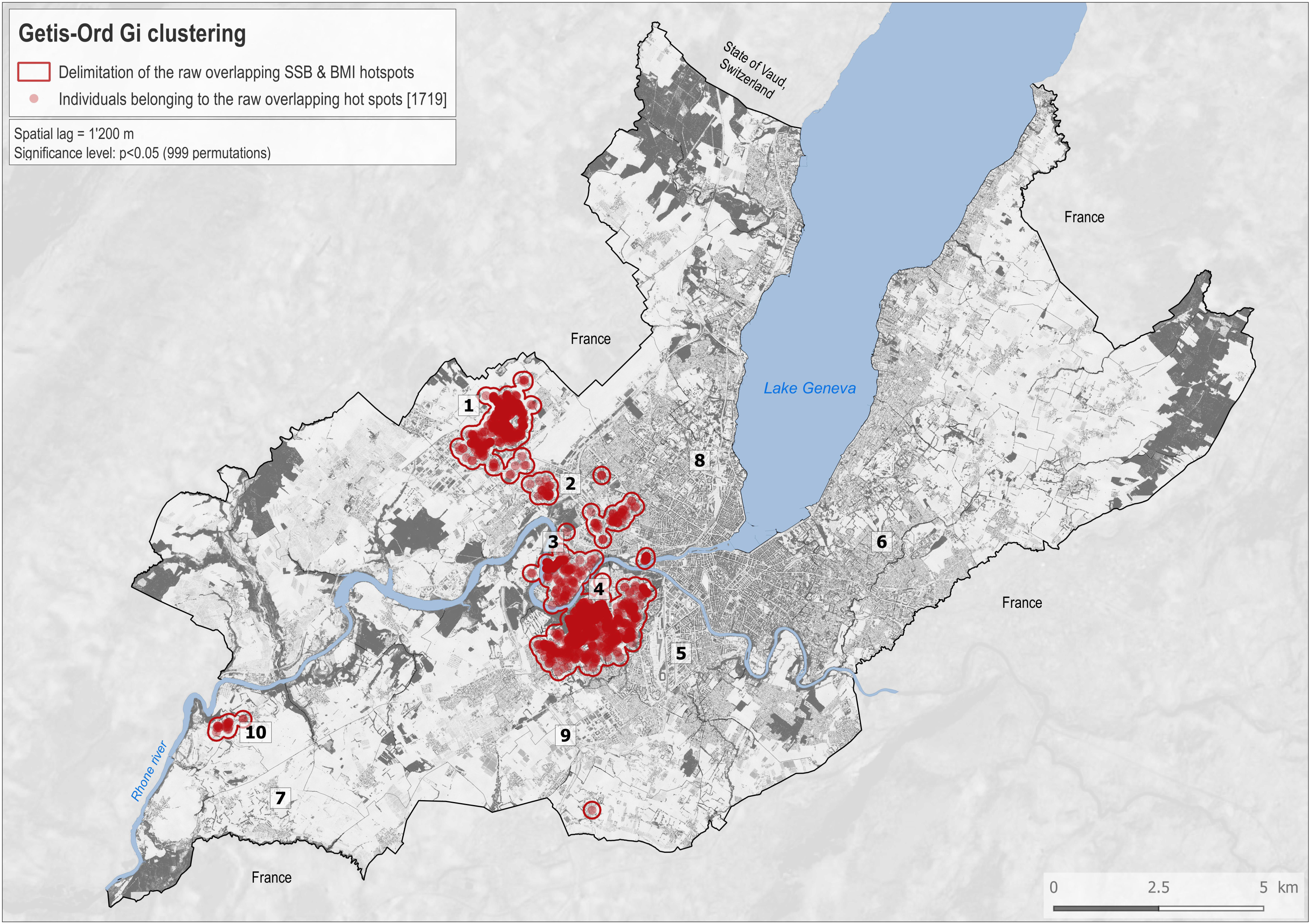
Maps showing the spatial distribution of local clusters of BMI and SSB intake frequency across subperiods P1 (A), P2 (B), and P3 (C).

**Figure S3:**
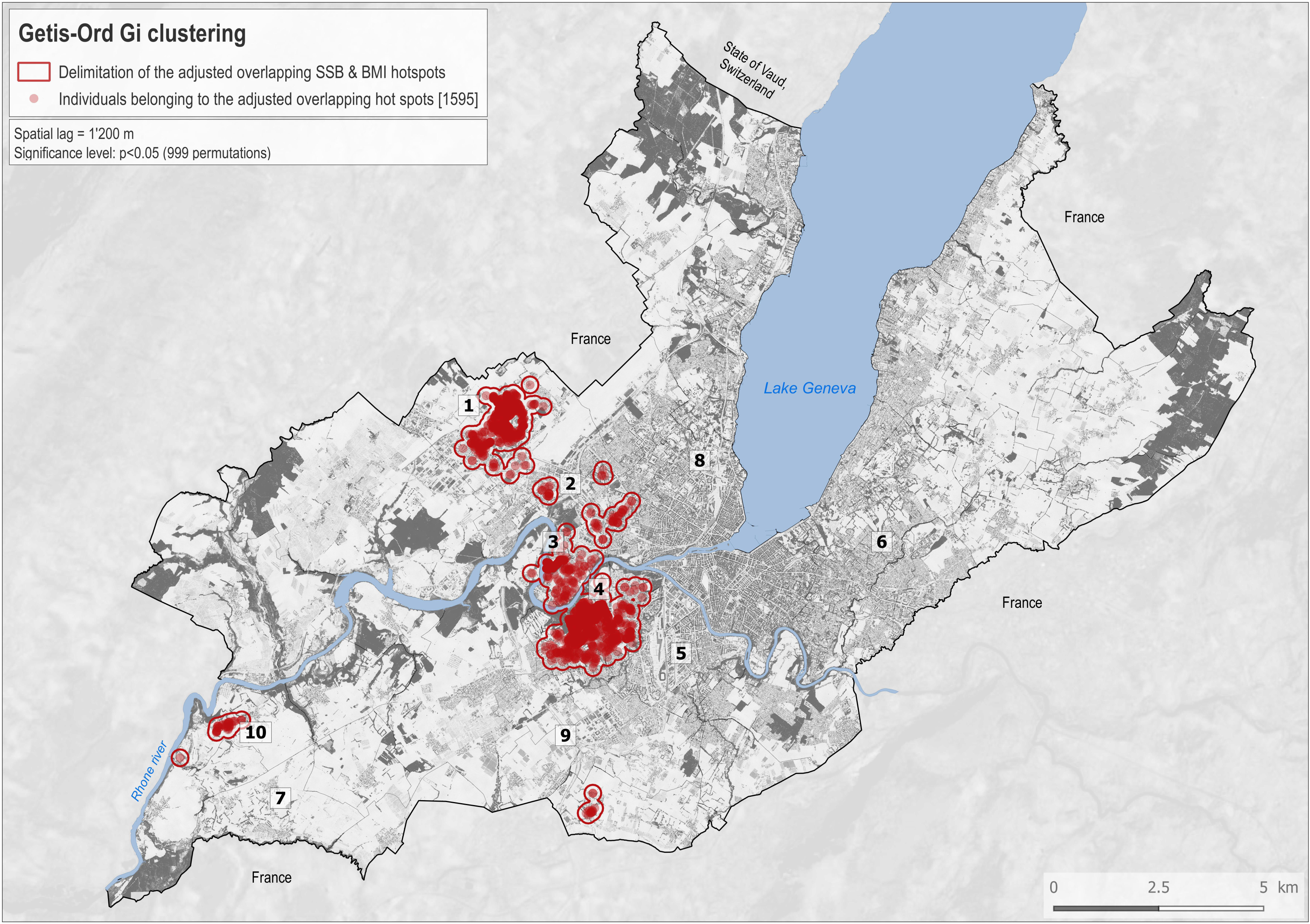
This map shows all the classes resulting from the combination of the different spatial autocorrelation regimes as measured by the Getis-Ord GI clustering for raw SSB intake frequency and raw BMI (A) and for adjusted SSB intake frequency and adjusted BMI (B). To facilitate reading, Figure 3A and 3B show only the overlapped hotspots for raw SSB intake frequency and raw BMI and adjusted SSB intake frequency and adjusted BMI, respectively.

**Figure S4:** Spatial distribution of the pseudo p-value for the adjusted BMI variable.

**Figure S5:** Spatial distribution of the pseudo p-value for the adjusted SSB variable.

## REFERENCES

1. Ritchie, Anne and Roser M. Obesity - Our World in Data [Internet]. 2018 [cited 2018 May 23]. Available from: https://ourworldindata.org/obesity

2. Ludwig DS, Peterson KE, Gortmaker SL. Relation between consumption of sugar-sweetened drinks and childhood obesity: a prospective, observational analysis. Lancet. 2001 Feb 17;357(9255):505–8.

3. Vereecken CA, Inchley J, Subramanian SV, Hublet A, Maes L. The relative influence of individual and contextual socio-economic status on consumption of fruit and soft drinks among adolescents in Europe. Eur J Public Health. 2005 Jun 1;15(3):224–32.

4. CDC. The CDC Guide to Strategies for Reducing the Consumption of Sugar-Sweetened Beverages.

5. Ventura EE, Davis JN, Goran MI. Sugar Content of Popular Sweetened Beverages Based on Objective Laboratory Analysis: Focus on Fructose Content. Obesity. 2011 Apr 14;19(4):868–74.

6. Market Research on the Soft Drinks Industry [Internet]. [cited 2018 Apr 9]. Available from: http://www.euromonitor.com/soft-drinks

7. Home - Unesda [Internet]. [cited 2018 Apr 9]. Available from: https://www.unesda.eu/

8. Promotion Santé Suisse Rapport 3.

9. Hill JO, Wyatt HR, Peters JC. Energy Balance and Obesity. Circulation. 2012 Jul 3;126(1):126–32.

10. Welsh JA, Lundeen EA, Stein AD. The sugar-sweetened beverage wars. Curr Opin Endocrinol Diabetes Obes. 2013 Oct;20(5):401–6.

11. Malik VS, Pan A, Willett WC, Hu FB. Sugar-sweetened beverages and weight gain in children and adults: a systematic review and meta-analysis. Am J Clin Nutr. 2013 Oct 1;98(4):1084–102.

12. Xi B, Huang Y, Reilly KH, Li S, Zheng R, Barrio-Lopez MT, et al. Sugar-sweetened beverages and risk of hypertension and CVD: a dose–response meta-analysis. Br J Nutr. 2015 Mar 4;113(05):709–17.

13. Cohen L, Curhan G, Forman J. Association of Sweetened Beverage Intake with Incident Hypertension. J Gen Intern Med. 2012 Sep 27;27(9):1127–34.

14. Larsson SC, Åkesson A, Wolk A. Sweetened Beverage Consumption Is Associated with Increased Risk of Stroke in Women and Men. J Nutr. 2014 Jun 1;144(6):856–60.

15. Bernstein AM, de Koning L, Flint AJ, Rexrode KM, Willett WC. Soda consumption and the risk of stroke in men and women. Am J Clin Nutr. 2012 May 1;95(5):1190–9.

16. Imamura F, O′Connor L, Ye Z, Mursu J, Hayashino Y, Bhupathiraju SN, et al. Consumption of sugar sweetened beverages, artificially sweetened beverages, and fruit juice and incidence of type 2 diabetes: systematic review, meta-analysis, and estimation of population attributable fraction. BMJ. 2015 Jul 21;351:h3576.

17. Vargas-Garcia EJ, Evans CEL, Prestwich A, Sykes-Muskett BJ, Hooson J, Cade JE. Interventions to reduce consumption of sugar-sweetened beverages or increase water intake: evidence from a systematic review and meta-analysis. Obes Rev. 2017 Nov;18(11):1350–63.

18. Brownell KD, Farley T, Willett WC, Popkin BM, Chaloupka FJ, Thompson JW, et al. The Public Health and Economic Benefits of Taxing Sugar-Sweetened Beverages. N Engl J Med. 2009 Oct 15;361(16):1599–605.

19. Auchincloss AH, Gebreab SY, Mair C, Diez Roux A V. A Review of Spatial Methods in Epidemiology, 2000–2010. Annu Rev Public Health. 2012 Apr 21;33(1):107–22.

20. Guessous I, Joost S, Jeannot E, Theler J-M, Mahler P, Gaspoz J-M, et al. A comparison of the spatial dependence of body mass index among adults and children in a Swiss general population. Nutr Diabetes. 2014 Mar 10;4(3):e111–e111.

21. Joost S, Haba-Rubio J, Himsl R, Vollenweider P, Preisig M, Waeber G, et al. Spatial clusters of daytime sleepiness and association with nighttime noise levels in a Swiss general population (GeoHypnoLaus). Int J Hyg Environ Health. 2018 Jun 1;

22. Park S, McGuire LC, Galuska DA. Regional Differences in Sugar-Sweetened Beverage Intake among US Adults. J Acad Nutr Diet. 2015 Dec;115(12):1996–2002.

23. Kumar GS, Pan L, Park S, Lee-Kwan SH, Onufrak S, Blanck HM, et al. Sugar-sweetened beverage consumption among adults -- 18 states, 2012. MMWR Morb Mortal Wkly Rep. 2014 Aug 15;63(32):686–90.

24. Han E, Powell LM. Consumption Patterns of Sugar-Sweetened Beverages in the United States. J Acad Nutr Diet. 2013 Jan;113(1):43–53.

25. Guessous I, Bochud M, Theler J-M, Gaspoz J-M, Pechère-Bertschi A. 1999–2009 Trends in Prevalence, Unawareness, Treatment and Control of Hypertension in Geneva, Switzerland. Barengo NC, editor. PLOS One. 2012 Jun 27;7(6):e39877.

26. Marques-Vidal P, Gaspoz JM, Theler JM, Guessous I. Twenty-year trends in dietary patterns in French-speaking Switzerland: Toward healthier eating. Am J Clin Nutr. 2017;106(1):217–24.

27. Bernstein M, Morabia A, Costanza MC, Landis JR, Ross A, Flandre P, et al. [Nutritional balance of the diet of the adult residents of Geneva]. Soz Praventivmed. 1994;39(6):333–44.

28. Beer-Borst S, Costanza MC, Pechère-Bertschi A, Morabia A. Twelve-year trends and correlates of dietary salt intakes for the general adult population of Geneva, Switzerland. Eur J Clin Nutr. 2009 Feb 10;63(2):155–64.

29. Mozaffarian D, Fahimi S, Singh GM, Micha R, Khatibzadeh S, Engell RE, et al. Global Sodium Consumption and Death from Cardiovascular Causes. N Engl J Med. 2014 Aug 14;371(7):624–34.

30. Micha R, Khatibzadeh S, Shi P, Fahimi S, Lim S, Andrews KG, et al. Global, regional, and national consumption levels of dietary fats and oils in 1990 and 2010: a systematic analysis including 266 country – specific nutrition surveys. BMJ. 2014 Apr 15;348:g2272.

31. Statistiques cantonales - République et canton de Genève [Internet]. [cited 2018 Apr 9]. Available from: https://www.ge.ch/statistique/

32. Getis A, Ord JK. The Analysis of Spatial Association by Use of Distance Statistics. Geogr Anal. 1992;24(3):189–206.

33. Ord JK, Getis A. Local Spatial Autocorrelation Statistics: Distributional Issues and an Application. Geogr Anal. 1995;27(4):286–306.

34. Anselin L, McCann M. OpenGeoDa, open source software for the exploration and visualization of geospatial data. In: Proceedings of the 17th ACM SIGSPATIAL International Conference on Advances in Geographic Information Systems - GIS ’09. New York, New York, USA: ACM Press; 2009. p. 550.

35. Anselin L. Local Indicators of Spatial Association-LISA. Geogr Anal. 2010 Sep 3;27(2):93–115.

36. Wei Y, Pere A, Koenker R, He X. Quantile regression methods for reference growth charts. Stat Med. 2006 Apr 30;25(8):1369–82.

37. Tukey JW. Comparing Individual Means in the Analysis of Variance. Source: Biometrics. 1949;5(2):99–114.

38. Singh GM, Micha R, Khatibzadeh S, Lim S, Ezzati M, Mozaffarian D, et al. Estimated Global, Regional, and National Disease Burdens Related to Sugar-Sweetened Beverage Consumption in 2010CLINICAL PERSPECTIVE. Circulation. 2015 Aug 25;132(8):639–66.

39. Chriqui JF, Chaloupka FJ, Powell LM, Eidson SS. A typology of beverage taxation: multiple approaches for obesity prevention and obesity prevention-related revenue generation. J Public Health Policy. 2013 Aug 23;34(3):403–23.

40. Marques-Vidal P, Rousi E, Paccaud F, Gaspoz J-M, Theler J-M, Bochud M, et al. Dietary Intake according to Gender and Education: A Twenty-Year Trend in a Swiss Adult Population. Nutrients. 2015 Nov 18;7(11):9558–72.

41. Bosma H, van de Mheen HD, Borsboom GJ, Mackenbach JP. Neighborhood socioeconomic status and all-cause mortality. Am J Epidemiol. 2001 Feb 15;153(4):363–71.

42. Christakis NA, Fowler JH. The Spread of Obesity in a Large Social Network over 32 Years. N Engl J Med. 2007;357(4):370–9.

43. Moodley G, Christofides N, Norris SA, Achia T, Hofman KJ. Obesogenic Environments: Access to and Advertising of Sugar-Sweetened Beverages in Soweto, South Africa, 2013. Prev Chronic Dis. 2015 Oct 29;12:140559.

44. Tamura K, Duncan DT, Athens JK, Bragg MA, Rienti M, Aldstadt J, et al. Geospatial clustering in sugar-sweetened beverage consumption among Boston youth. Int J Food Sci Nutr. 2017 Aug 18;68(6):719–25.

45. Lesser LI, Zimmerman FJ, Cohen DA. Outdoor advertising, obesity, and soda consumption: a cross-sectional study. BMC Public Health. 2013 Jan 10;13:20.

46. Wiecha JL, Finkelstein D, Troped PJ, Fragala M, Peterson KE. School Vending Machine Use and Fast-Food Restaurant Use Are Associated with Sugar-Sweetened Beverage Intake in Youth. J Am Diet Assoc. 2006 Oct;106(10):1624–30.

47. Stanhope KL. Sugar consumption, metabolic disease and obesity: The state of the controversy. Crit Rev Clin Lab Sci. 2016 Jan 2;53(1):52–67.

